# Flooding tolerance of four tropical peatland tree species in a nursery trial

**DOI:** 10.1101/2021.12.26.474202

**Authors:** Hesti L. Tata, Hani S. Nuroniah, Diandra A. Ahsania, Haning Anggunira, Siti N. Hidayati, Meydina Pratama, Istomo, Rodney A. Chimner, Meine van Noordwijk, Randall Kolka

## Abstract

In order to facilitate hydrological restoration efforts, initiatives have been conducted to promote tree growth in degraded and rewetted peatlands in Indonesia. For these initiatives to be successful, tree seedlings need to be able to survive flooding episodes, with or without shade. We investigated the survival rates and the formation of adventitious roots in the case of four tree species exposed to combinations of different shading and water levels under controlled conditions in a nursery, with artificial rainwater and with peat soils as the medium. The research focused on the following questions (i) whether trees can grow on flooded peat soils; and (ii) which plant traits allow plants to cope with inundation, with or without shade. The four tree species compared (*Shorea balangeran, Cratoxylum arborescens, Nephelium lappaceum* and Durio *zibethinus*) include two natural pioneer and two farmer-preferred fruit trees. The experiment used a split-split plot design with 48 treatment combinations and at least 13 tree-level replicates. The study found that *S. balangeran* and C. *arborescens* had relatively high survival rates and tolerated saturated condition for 13 weeks, while *N. lappaceum* and *D. zibethinus* required non-saturated peat conditions. *S. balangeran* and C. *arborescens* developed adventitious roots to adapt to the inundated conditions. *D. zibethinus, S. balangeran* and *N. lappaceum* grew best under moderate (30%) shading levels, while *C. arborescent* grew best in full sunlight.

## Introduction

Tropical peatland ecosystems [1] harbor globally unique biodiversity, regulating water flows and storing carbon. Global peatlands store about 50 –180 Pg of carbon [2] that, if released into the atmosphere, would be equivalent to 6 – 22 years of current global fossil fuel emissions. In their natural condition, peatlands are saturated most of the time, with a distinct microtopography providing drier regeneration niches [3].

Wetland vegetation has adapted to thrive in anerobic conditions, characterized by difficult oxygen supply to roots, through the application of a number of adaptation strategies, including the development of aerenchyma tissue, lenticels, adventitious roots and pneumatophores [4–9]. The limited availability of soil nutrients favors plants that have insectivorous nutrition or nutrient-conserving properties, such as thick leaves [10–11].

To facilitate the restoration of degraded Indonesian peatlands, canals that were initially built to transport logs have been replaced by deep channels to drain peat and to enable the conversion of the land for agricultural use. This drainage process accelerates peat oxidation, increasing CO_2_ emissions and leading to subsidence, consequently increasing susceptibility to drought and fire and exacerbating downstream flooding risks [12,13,14]. On drained peatlands, farmers cultivate non-peatland species that are not adapted to wet condition.

The issue of peatland restoration has become critically important in Indonesia, particularly after extreme fire events in 2015 that affected more than 2.6 Mha of forest lands, including 0.89 Mha of peatlands [15]. In Indonesia, successful peatland restoration initiatives often involve interventions such as hydrological restoration, revegetation using suitable plant species, and measures to improve the livelihoods of local communities. In particular, hydrological restoration involves canal blocking and rewetting. These activities are intended to increase water levels and peat humidity [16,17]. While more than 500 plant species grow naturally on peatlands [18], most of these wetland species have low economic value compared to the non-peatland tree species with which they are often replaced, particularly sengon (*Falcataria moluccana*) [19] and oil palm (*Elaeis guinensis*) [20]. While the economic benefits generated from cultivating these introduced tree species planted on peatlands is significant, little consideration is given to the suitability of these species under these conditions [21].

To facilitate the restoration of peatlands, canals or diches are blocked to increase groundwater levels [22–24] and thereby to reduce susceptibility to fire [23]. The intervention is also intended to restore the function of the peat as a carbon sink and to reduce carbon emissions [25]. Direct barriers to peatland restoration initiatives in Indonesia include altered peat topography, peatland drainage, the presence of invasive ferns and shrub species, fires, and flooding risks, while indirect barriers include climate change, inconsistent land-use policy, and lack of sustainable livelihood options for members of communities in surrounding areas [26]. The natural regeneration of peat-swamp forest on degraded peatlands occurs as a result of mammal and bird dispersal and is correlated with the distance of the site to the forest, with regeneration occurring mostly in clusters of young trees at distances of up to 2 km from the nearest forest [26]. At further distances, it may be necessary to plant trees to facilitate the establishment process. However, it can be shown that current peatland restoration practices do not achieve the aim of reducing carbon emissions due to the poor design of interventions, with only some of the existing canals being blocked (which means that water levels do not rise to a sufficient level), or with unsuitable tree species being planted for paludiculture [27,28].

In order to restore peatlands where the increases in the water levels or the influx of natural seeds are insufficient, trees that are adapted to wet condition should be planted. The application of paludiculture can therefore serve as an alternative means of overcoming the constraints on mitigating carbon emissions through peatland restoration initiatives, so long as the trees utilized for the revegetation program are suitable to the wet (and rewetted) conditions that prevail within the restored peatlands [19,29,30].

In the case of Indonesia’s peatland restoration program, increasing the ground water level is a precondition for the success of the initiative [16]. The Minister of Environment and Forestry Regulation No. 16, 2017, includes a list of plant species that have been deemed suitable for peatland restoration. However, limited information related to the root systems of tropical plants in saturated conditions is available. The first experimental study into the root systems of 65 tropical tree species in flooded conditions was conducted in 1932, with the study finding that *Glutha renghas* was able to survive in such conditions for a period of up to a year [32]. In a more recent study, 26 native peatland tree species were investigated to determine their level of tolerance to shading and flooding. It was found that *Lophopetalum javanicum* had the best survival rate in flooded conditions, with two fruit tree species (*Mangifera* sp. and *Syzygium* sp.) categorized as *light-demanding* species [33]. The level of mycorrhizal dependence of *S. balangeran* was also investigated in the context of peatland restoration initiatives [34].

A number of different tree species have been planted to facilitate the restoration of peatlands, including timber tree species that do not grow naturally swamp forests, particularly *Shorea balangeran* [24,35] and *Cratoxylum arborescens* [36,37]. In the timber trade, *S. balangeran* (in the Dipterocarpaceae family) is known as ‘red meranti,’ with timber from this species being highly durable and thus utilized for the construction of bridges, housing, and a wide range of other purposes [38,39]. *C. arboresencens* yields a lightweight timber with long fibers, which is thus suitable as a raw material for the pulp and paper industry [40] and for use in light construction, including the production of plywood, transportation boxes, and utensils [38,41].

In addition, a range of fruit tree species, including *Nephelium lappaceum* [28,42] (and *Durio zibethinus* [37,43,44], have also been planted as part of peatland restoration initiatives, particularly in agroforestry systems, where they are planted together with other tree species. While *N. lappaceum* and *D. zibethinus* are not naturally distributed in peat swamp forest [45,46], these two fruit tree species are commonly planted on peatlands by local communities in Central Kalimantan because they have high economic value, with strong demand from local and national markets [42,43].

However, there is a lack of information related to the conditions under which four commonly planted species should be utilized in these rewetted peatlands in Central Kalimantan. To address this, this study aims to compare the growth of two timber tree-species (*C. arborescens* and *S. balangeran*) and two fruit tree species (*D. zibethinus* and *N. lappaceum*) in varying conditions of inundation and shading through a controlled mesocosm study in a nursery, under a range of different water table levels and light conditions. The inundation treatment was designed to simulate the saturated conditions found in rewetted peatlands, while the shading treatment was intended to test levels of tolerance to direct sunlight.

## Materials and methods

### Study site

The study was conducted in a nursery managed by the Forest Research and Development Center in Bogor, Indonesia. Peat soils were collected from Kameloh village, in Palangkaraya, Central Kalimantan. The planting stocks used for the experiment were collected from two locations: two peatland tree-species, *C. arborescens* and *S. balangeran*, were collected from wildings in Tumbang Nusa, Central Kalimantan, while the two fruit tree species, *D. zibethinus* and *N. lappaceum*, were germinated from seeds collected in Bogor, West Java. The seedlings were planted in polyethylene bags (polybag) at 20 cm in height.

### Experimental design

In the nursery, the study involved the use of a plastic box container, which enabled the saturation level to be Tombing adjusted manually (a simple mesocosm study). A mesocosm study is usually conducted to simulate environmental parameters, using sophisticated equipment and automated procedures [47,48]. This experiment used a split-split plot design, with tree species in the main plot, with the shading treatment conducted in the sub-plot, and the inundation treatment conducted in the sub-sub-plot. It involved four different types of tree species: *C. arboresencens, S. balangeran, D. zibethinus* and *N. lappaceum*. The shading treatment involved three levels, at 30%, 70% and control (without shading). The flooding or inundation treatment involved four levels of control: freely drained; ground water (GW) level 10 cm (half of polybag’s height); GW level zero (equal to the polybag’s height); and ⅓ stem height flooding (i.e., 9 cm for *C. arborescens*, 8 cm for *S. balangeran*, 10 cm for *D. zibethinus* and 7 cm for *N. lappaceum)*. Due to uneven germination, the replication number four each treatment combination varied among plant species, with 13 seedlings for *C. arborescens*; 15 seedlings for *D. zibethinus* and *N. lappaceum*; and 20 seedlings for *S. balangeran*. The total number of seedlings of the *C. arborescens, D. zibethinus, N. lappaceum* and *S. balangeran* species stood at 156, 180, 180 and 240, respectively.

Each treatment combination was applied in a plastic container measuring 30×30×50 cm^3^. Two different parafilm-plastic-nets were utilized for the shading treatment, with a 70% and 30% light transmission rate respectively. The samples subject to shading were placed in a screen house, with the non-shaded control placed in an open area nearby. The layout of the treatment combination is shown in Fig 1.

**Fig 1.**
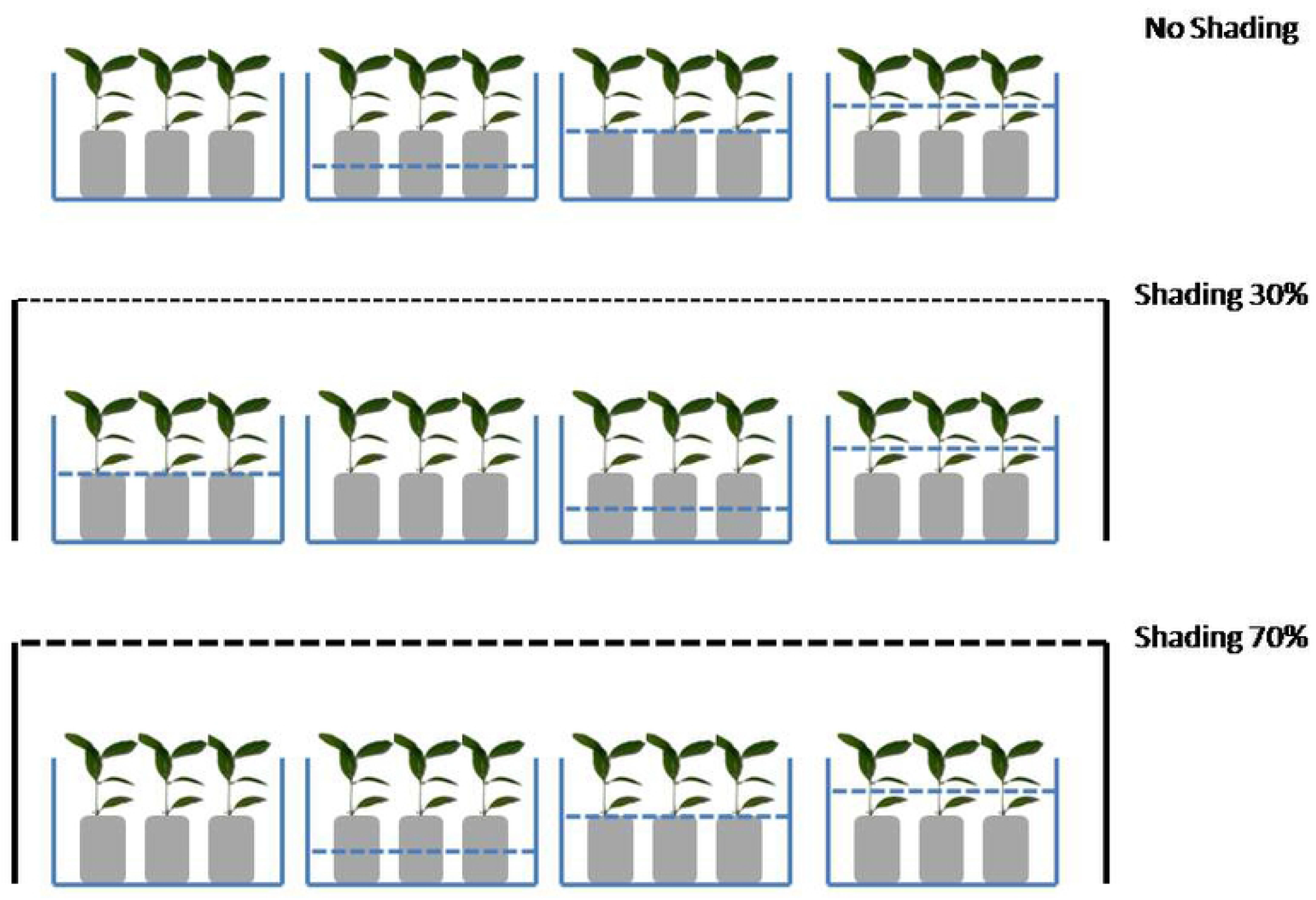
Layout of shading and inundation experiment in the nursery. Blue dashes indicate the ground water (GW) level, at GW level 10 cm (half of polybag’s height); GW level zero (the height of the polybag), and ⅓ stem height flooding. The diagram with no blue dashes indicates the control treatment (fully drained).

### Artificial rainwater

Artificial rainwater was created according to a formula that replicated the chemical properties of rainwater [49]. This water was poured into the plastic containers to meet the respective inundation levels. The control (non-saturated) seedlings were watered everyday with 10 ml rainwater. The containers were placed randomly in the space available at every shading level.

### Observation and growth measurement

A number of variables were observed, including survival rate, growth of stem diameter, growth of height, photosynthetic rate, chlorophyll content, and potential redox of the rainwater in the container, throughout the thirteen weeks during which the experiment was conducted. Measurements were taken once a week for the 13 weeks period.

To determine the diameter of the stem of the plant samples, measurements were taken at the base of the stem, about 1 cm above the soil media. The position of the growth measurement was marked using a permanent marker prior to the experiment. Stem diameter and total height was measured using a caliper and ruler, respectively. Measurements were taken once a week for the 13-weeks period.

The dry weights of shoots and roots were measured during the last week of the experiment. Three samples of surviving seedlings were selected purposively on the basis of their height and harvested to enable measurement of their dry weight. At harvest, the roots were cleaned carefully to remove soil. All part of the seedlings were dried and weighed. The shoot component consisted of leaves, stem, and aboveground adventitious roots that grow on the stem; the root component consisted of the roots that were submerged in the soil. The ratio of root and shoot weight is an important index for assessing plant health under stress condition [50,51]. Thus, we calculated the root/shoot ratio of the seedings based on the dry weight of the seedling samples.

The photosynthetic rate of each treatment was measured using a portable photosynthesis system, LI-COR 6400XT (LI-COR Inc., Lincoln, Nebraska), with this measurement taking place once a month for three months. Three leaves of leaf number 1, 2 and 3 from the shoot were measured. The photosynthetic rate was measured for each of the different shading levels. The LI-COR device was set at the following Photosynthetically Active Radiation (PAR) levels, depending on the respective shading level for each of the treatment samples (70%, 30% and control, 0%) at 100, 150 and 1000 µmol m^-2^ sec^-1^, respectively. The measurement was conducted in the morning at between 09:00 to 11:00 A.M.

Many peatland plant species are characterized by morphological adaptations, including adventitious roots formation, that enable them to grow in saturated soil conditions [52]. Adventitious roots are defined as roots originating from shoot tissue [53]. We observed and counted adventitious roots (AR) from all the seedlings for the four plants species tested every week.

Prior to the experiment, a sub-sample of peat soil was sent to the laboratory of Soil and Land Resources Department at the IPB University in Bogor to analyze its soil chemical properties, including soil pH, C-organic content, N-total, P-available, cation exchange capacity (CEC), pyrite concentration, ash content, and fiber content. In addition, abiotic factors were also measured three times daily throughout the three-month period for all of the three shading levels. Daily light intensity was measured using a Lux meter, while air temperature and humidity were measured using a thermo-hygrometer.

### Data analysis

The growth and survival of seedlings were measured every week for 13 weeks. Owing to the high mortality rate for the *D. zibethinus* and *N. lappaceum* seedlings, the growth of height and diameter for these species were measured up to 9 weeks only. The relative growth rate (RGR) of diameter and height was measured using the following formula [54]:

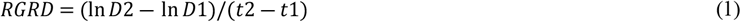

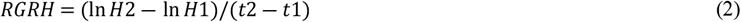

Where: RGRH: relative growth rate of height

RGRD: relative growth rate of diameter

D1, H1: Diameter, height of first measurement

D2, H2: Diameter or height of second measurement

T1 : first measurement

T2 : ninth measurement (at \9 weeks after planting)

Adventitious roots (AR) were found in the case of two species, *C. cratoxylum* and *S. balangeran*. The AR formation of all *C. arborescens* seedlings was analyzed at 9 weeks after planting, while the AR formation of *S. balangeran* was analyzed on the basis of a sub-sample of 3 seedlings at 13 weeks after planting.

The variables related to the survival rate, relative growth rate of diameter (RGRD), relative growth rate of height (RGRH), dry weight of shoot, dry weight of roots, root/shoot ratio and number of adventitious roots were analyzed using a General Linear Model (GLM) in a univariate analysis. The Duncan Multiple Range Test (DMRT) was applied as post-hoc test when *p*-value < 0.05.

The photosynthetic rate of four plant species was analyzed based on an analysis of covariance, with Photosynthetically Active Radiation (PAR) at 3 shading levels used as covariates. SPSS ver.21 for IBM was used to conduct the data analysis.

In addition, the distribution of the dry weight of the shoot and root of each species at different inundation level was analyzed by conducting a regression analysis for each species to determine the allometric equation for each species.

## Results

### Survival rate

An examination of the weekly survival rate for each of the four species used in the experiment showed that *S. balangeran* had the highest survival rate, while the *D. zibethinus* had the lowest. *C. arborescens* did not survive in the combination treatment of saturation at ⅓ stem height (9 cm flooding) and 70% shading (Fig 2).

**Fig 2.**
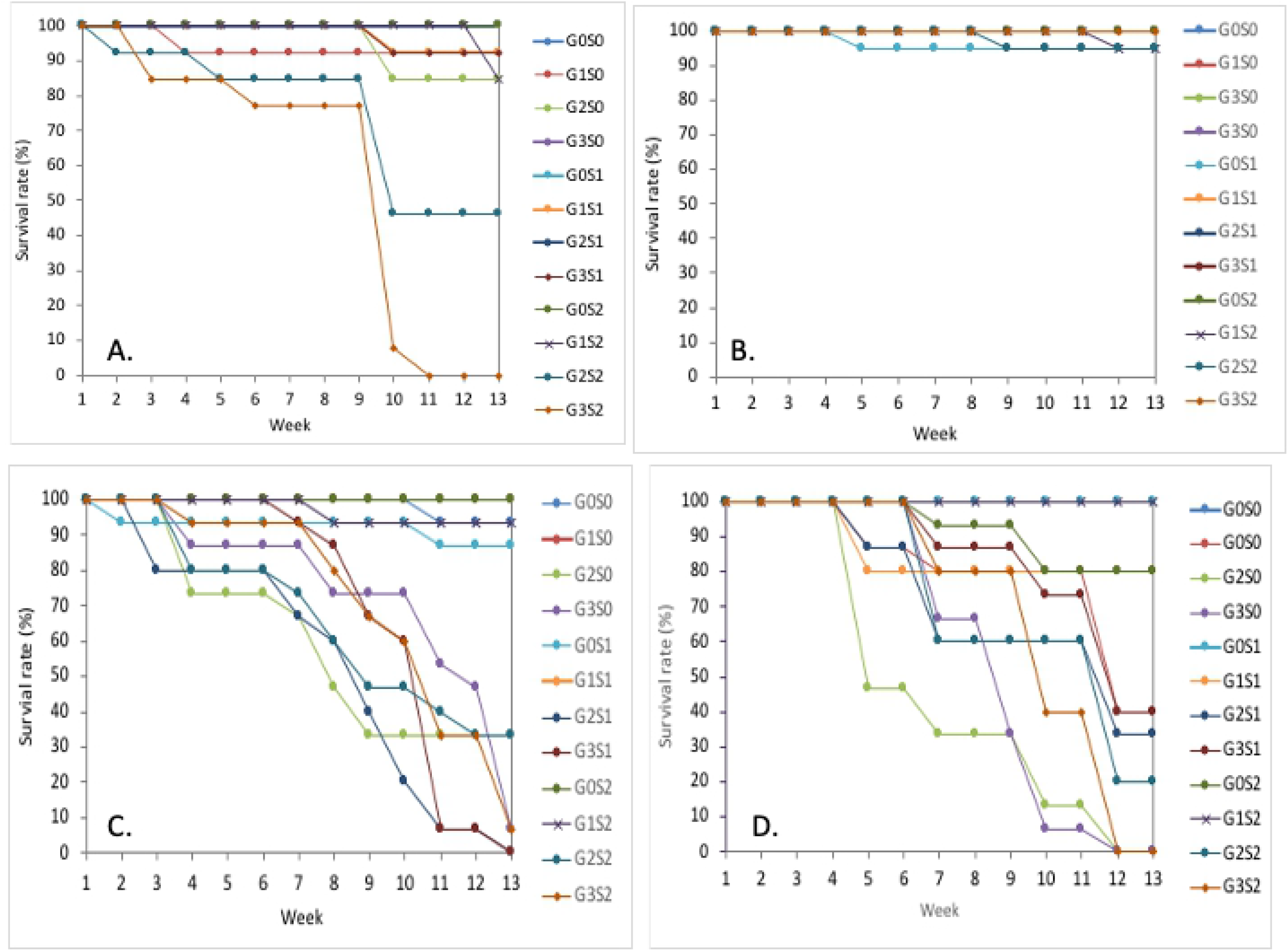
Survival rate of four tropical peatland tree species. A) *Cratoxylum arborescens*, b) *Shorea balangeran*, c) *Nephelium lappaceum*, d) *Durio zibethinus*, in treatment combination of shading and saturated condition. G0S0= Freely drained without shading, G0S1= Freely drained with 30% shading, G0S2= Freely drained with 70% shading, G1S0 = GW level 10 cm without shading, G1S1= GW level 10 cm with 30% shading, G1S2= GW level 10 cm with 70% shading, G2S0= GW level zero without shading, G2S1= GW level zero with 30% shading, G2S2= GW level zero with 70% shading, G3S0= ⅓ stem height flooding without shading, G3S1= ⅓ stem height flooding with 30% shading, G3S2= ⅓ stem height flooding with 70% shading. A statistical analysis of the interaction between plant species, inundation and shading levels (Table 1) confirmed the differences in the GLM analysis for each treatment and their interaction effect to the survival rate, with the results of this analysis shown in the following S1 Table.

The four plants species grew well in the freely drained soil, despite the fact that *C. arborescens* and *S. balangeran* are peatland tree species. The analysis of the soil samples prior to the experiment (S6 Table) showed that the soil pH was low (pH KCl=3.18); the C organic content was 54.96%; the Nitrogen content was 0.55% (middle); and the ash content was 5.24%. The soil sample was collected from degraded peatland and is categorized as hemic sapric. The peat soils support the growth of the seedlings of the four plants species in the control treatment (Fig 2, Table 1 and 2).

**Table 1.** Survival rate of four plant species under different treatment.

| Treatment | Survival rate (%) |  |  |  |
| --- | --- | --- | --- | --- |
|  | *C. arborescens* | *S. balangeran* | *D. zibethinus* | *N. lappaceum* |
| Freely drained without shading | 100.0 ± 0.0 a | 100.0 ± 0.0 a | 100.0 ± 0.0 a | 98.5 ± 2.9 ab |
| Freely drained with 30% shading | 100.0 ± 0.0 a | 96.5 ± 2.4 b | 100.0 ± 0.0 a | 92.3 ± 3.7 b |
| Freely drained with 70% shading | 100.0 ± 0.0 a | 100.0 ± 0.0 a | 92.3 ± 9.0 b | 100.0 ± 0.0 a |
| GW level 10 cm without shading | 94.1 ± 3.4 bc | 100.0 ± 0.0 a | 81.0 ± 20.2 bc | 100.0 ± 0.0 a |
| GW level 10 cm with 30% shading | 97.6 ± 3.7 ab | 100.0 ± 0.0 a | 86.2 ± 9.6 bc | 100.0 ± 0.0 a |
| GW level 10 cm with 70% shading | 98.8 ± 4.3 ab | 99.2 ± 1.8 ab | 100.0 ± 0.0 a | 96.9 ± 3.5 b |
| GW level zero without shading | 95.3 ± 7.4 bc | 100.0 ± 0.0 a | 47.7 ± 39.2 d | 61.5 ± 27.5 cd |
| GW level zero with 30% shading | 100.0 ± 0.0 a | 100.0 ± 0.0 a | 72.3 ± 24.6 bc | 55.4 ± 36.3 cd |
| GW level zero with 70% shading | 75.7 ± 21.0 bc | 98.1 ± 2.5 b | 72.3 ± 30.0 bc | 67.2 ± 25.3 cd |
| 1/3 stem height flooding without shading | 100.0 ± 0.0 a | 100.0 ± 0.0 a | 60.0 ± 44.1 c | 74.9 ± 26.4 bc |
| 1/3 stem height flooding with 30% shading | 97.6 ± 3.7 b | 100.0 ± 0.0 a | 83.6 ± 21.7 b | 70.8 ± 40.0 bc |
| 1/3 stem height flooding with 70% shading | 59.2 ± 40.5 c | 100.0 ± 0.0 a | 70.8 ± 37.9 bc | 73.3 ± 31.2 bc |
| Grand mean (SEm) | 93.2 (1.4) g | 99.5 ± (1.4) f | 80.5 (1.4) h | 82.5 (1.4) h |

Note: Means are followed by standard deviation. Value in the same column followed by different letter is significantly different at $p$ -value <0.05. Grand mean is compared among species. S.E.m is standard error of mean.

**Table 2.** RGRD of *C. arborescens, S. balangeran, D. zibethinus* and *N. lappaceum* at 9 weeks after planting in the shading and inundation treatments.

| Treatment | RGRD at 9 weeks |  |  |  |
| --- | --- | --- | --- | --- |
|  | *C. arborescens* | *S. balangeran* | *D. zibethinus* | *N. lappaceum* |
| Freely drained without shading | 0.051 ± 0.027 b | 0.072 ± 0.029 a | 0.023 ± 0.013 c | 0.063 ± 0.016 ab |
| Freely drained with 30% shading | 0.064 ± 0.015 a | 0.058 ± 0.015 abc | 0.018 ± 0.005 cd | 0.061 ± 0.021 ab |
| Freely drained with 70% shading | 0.024 ± 0.013 cd | 0.047 ± 0.016 bc | 0.015 ± 0.007 d | 0.036 ± 0.011 c |
| GW level 10 cm without shading | 0.054 ± 0.022 bc | 0.052 ± 0.020 abc | 0.015 ± 0.009 d | 0.070 ± 0.014 a |
| GW level 10 cm with 30% shading | 0.042 ± 0.013 bc | 0.048 ± 0.021 bc | 0.019 ± 0.010 cd | 0.061 ± 0.010 ab |
| GW level 10 cm with 70% shading | 0.023 ± 0.008 cd | 0.032 ± 0.016 c | 0.021 ± 0.009 c | 0.036 ± 0.016 c |
| GW level zero without shading | 0.056 ± 0.018 abc | 0.070 ± 0.019 ab | 0.009 ± 0.004 e | 0.055 ± 0.014 bc |
| GW level zero with 30% shading | 0.068 ± 0.009 a | 0.049 ± 0.016 bc | 0.018 ± 0.007 cd | 0.026 ± 0.012 cd |
| GW level zero with 70% shading | 0.014 ± 0.010 d | 0.040 ± 0.021 bc | 0.010 ± 0.007 de | 0.030 ± 0.003 cd |
| 1/3 stem height flooding without shading | 0.061 ± 0.012 ab | 0.049 ± 0.020 bc | 0.029 ± 0.0 c | 0.027 ± 0.015cd |
| 1/3 stem height flooding with 30% shading | 0.037 ± 0.015 c | 0.036 ± 0.026 bc | 0.012 ± 0.005 de | 0.032 ± 0.019 c |
| 1/3 stem height flooding with 70% shading | 0.013 ± 0.009 d | 0.033 ± 0.022 c | 0.007 ± 0.010 e | 0.017 ± 0.013 d |
| Grand Mean (S.E.m) | 0.042 (0.001) g | 0.055 (0.001) f | 0.016 (0.002) h | 0.043 (0.002) g |

Note: Means are followed by standard deviation. Value in the same column followed by different letter is significantly different at $p$ -value <0.05. Grand mean is compared among species. S.E.m. is a standard error of mean.

#### Relative Growth Rate of Diameter and Height (RGRD and RGRH)

The growth of diameter and height were measured once each week throughout the 13-week period. However, due to the high mortality rate for *D. zibethinus* and *N. lappaceum* at 10 weeks (Fig 2c and 2d), the RGRD and RGRH were then calculated and analyzed for the seedlings at 9 weeks after planting. The RGRD and RGRH are shown in Table 2 and 3, respectively. The GLM analysis showed that treatments of plant species, inundation levels, shading and their interactions affected the RGRD and RGRH significantly (S2 Table, S3 Table).

**Table 3.**
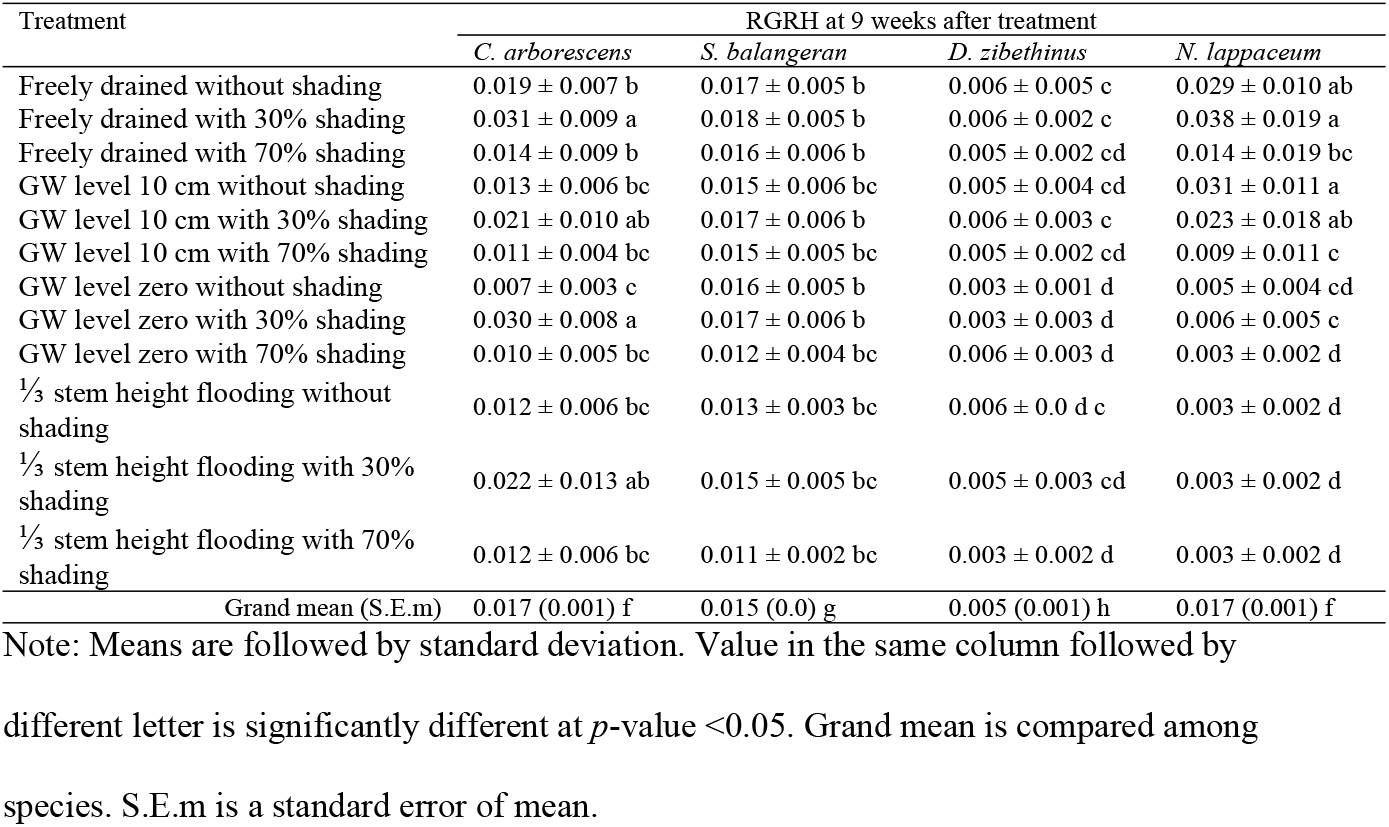
RGRH of *C. arborescens, S. balangeran, D. zibethinus* and *N. lappaceum* at 9 weeks after planting in the shading and inundation treatments.

| Treatment | RGRH at 9 weeks after treatment |  |  |  |
| --- | --- | --- | --- | --- |
|  | *C. arborescens* | *S. balangeran* | *D. zibethinus* | *N. lappaceum* |
| Freely drained without shading | 0.019 ± 0.007 b | 0.017 ± 0.005 b | 0.006 ± 0.005 c | 0.029 ± 0.010 ab |
| Freely drained with 30% shading | 0.031 ± 0.009 a | 0.018 ± 0.005 b | 0.006 ± 0.002 c | 0.038 ± 0.019 a |
| Freely drained with 70% shading | 0.014 ± 0.009 b | 0.016 ± 0.006 b | 0.005 ± 0.002 cd | 0.014 ± 0.019 bc |
| GW level 10 cm without shading | 0.013 ± 0.006 bc | 0.015 ± 0.006 bc | 0.005 ± 0.004 cd | 0.031 ± 0.011 a |
| GW level 10 cm with 30% shading | 0.021 ± 0.010 ab | 0.017 ± 0.006 b | 0.006 ± 0.003 c | 0.023 ± 0.018 ab |
| GW level 10 cm with 70% shading | 0.011 ± 0.004 bc | 0.015 ± 0.005 bc | 0.005 ± 0.002 cd | 0.009 ± 0.011 c |
| GW level zero without shading | 0.007 ± 0.003 c | 0.016 ± 0.005 b | 0.003 ± 0.001 d | 0.005 ± 0.004 cd |
| GW level zero with 30% shading | 0.030 ± 0.008 a | 0.017 ± 0.006 b | 0.003 ± 0.003 d | 0.006 ± 0.005 c |
| GW level zero with 70% shading | 0.010 ± 0.005 bc | 0.012 ± 0.004 bc | 0.006 ± 0.003 d | 0.003 ± 0.002 d |
| 1/3 stem height flooding without shading | 0.012 ± 0.006 bc | 0.013 ± 0.003 bc | 0.006 ± 0.0 d c | 0.003 ± 0.002 d |
| 1/3 stem height flooding with 30% shading | 0.022 ± 0.013 ab | 0.015 ± 0.005 bc | 0.005 ± 0.003 cd | 0.003 ± 0.002 d |
| 1/3 stem height flooding with 70% shading | 0.012 ± 0.006 bc | 0.011 ± 0.002 bc | 0.003 ± 0.002 d | 0.003 ± 0.002 d |
| Grand mean (S.E.m) | 0.017 (0.001) f | 0.015 (0.0) g | 0.005 (0.001) h | 0.017 (0.001) f |

Note: Means are followed by standard deviation. Value in the same column followed by different letter is significantly different at $p$ -value <0.05. Grand mean is compared among species. S.E.m is a standard error of mean.

Three plant species (*S. balangeran, D. zibethninus* and *N. lappaceum*) recorded the best RGRD results in the control treatment (unsaturated and without shading). Among the four species, *S. balangeran* had the highest RGRD, while *D. zibethinus* had the lowest RGRD (Table 2).

The best RGRH results were found in the case of the unsaturated treatment with 30% shading. Among all four plants species, *C. arborescens* had the highest RGRH. The treatment with ⅓ stem height flooding with 70% shading resulted in the worst RGRH for all of the four plant species (Table 3).

#### Dry weight of shoot, root and root/shoot ratio

A General Linear Model (GLM) analysis showed that the dry weight (DW) of the shoot, root, and S/R ratios were affected by all the treatment combinations (S4 Table). Among all four species, the dry weight of the shoots was highest in the case of *D. zibethinus*. The results for the dried weight of shoots varied between the species. In the case of *C. arborescens*, the results for the dry weight of the shoots were consistent, 30% shading resulted in the highest dry shoot weights at every inundation level. In the case of the other three species, *S. balangeran, D. zibethinus* and *N. lappaceum*, the results varied between the unsaturated and high-water level treatments. At inundation, the dry weight of shoot decreased with the increase of shading level. However, in the high-water-level treatments, the highest dry shoot weight resulted from 30% shading, with this rate decreasing at 70% shading (Table 4).

**Table 4.** Dry weight (DW) of shoots of four plant species under treatment combination of shading and inundation.

| Treatment | Dry weight of shoots (g) |  |  |  |
| --- | --- | --- | --- | --- |
|  | *C. arborescens* | *S. balangeran* | *D. zibethinus* | *N. lappaceum* |
| Freely drained without shading | 0.81 ± 0.05 cd | 0.69 ± 0.14 d | 4.61 ± 1.26 a | 2.13 ± 0.90 bc |
| Freely drained with 30% shading | 0.85 ± 0.21 cd | 0.67 ± 0.05 d | 2.17 ± 1.07 bc | 1.25 ± 0.14 cd |
| Freely drained with 70% shading | 0.36 ± 0.13 ef | 0.38 ± 0.05 de | 1.31 ± 0.21 c | 0.77 ± 0.43 cd |
| GW level 10 cm without shading | 0.89 ± 0.36 cd | 0.74 ± 0.03 cd | 3.17 ± 0.28 ab | 1.25 ± 0.28 cd |
| GW level 10 cm with 30% shading | 1.14 ± 0.14 cd | 0.88 ± 0.28 cd | 3.17 ± 0.32 ab | 1.92 ± 0.41 bc |
| GW level 10 cm with 70% shading | 0.68 ± 0.10 d | 0.37 ± 0.07 e | 2.34 ± 0.63 bc | 0.82 ± 0.20 cd |
| GW level zero without shading | 1.39 ± 0.44 cd | 0.73 ± 0.17 bc | - | 0.79 ± 0.45 cd |
| GW level zero with 30% shading | 1.55 ± 0.21 c | 0.61 ± 0.09 d | 2.98 ± 0.38 b | - |
| GW level zero with 70% shading | 0.45 ± 0.12 de | 0.35 ± 0.13 e | 1.18 ± 1.33 bc | 0.35 ± 0.30 def |
| 1/3 stem height flooding without shading | 1.25 ± 0.36 cd | 0.95 ± 0.02 cd | - | 0.22 ± 0.38 ef |
| 1/3 stem height flooding with 30% shading | 1.13 ± 0.17 cd | 1.16 ± 0.31 cd | 1.66 ± 0.42 bc | - |
| 1/3 stem height flooding with 70% shading | - | 0.44 ± 0.13 de | - | - |
| Grand mean (S.E.m) | 0.88 (0.22) n | 0.66 (0.22) n | 1.98 (0.22) m | 0.82 (0.22) n |

Note: Blank (-) indicates seedlings' mortality. A value followed by a different letter is
significantly different at $p < 0.05$ . Grand mean is compared among species, with the value in parentheses a standard error of mean.

In the control treatment (unsaturated without shading), the four plants species recorded the highest dry shoot weights. Under unsaturated conditions, the dry root weight decreased with increasing shading level. Among the four plants species, *C. arborescens* and *D. zibethinus* recorded the highest dry root weights. The response of the various plant species varied between treatments. In the case of *S. balangeran*, the dry weight consistently decreased with increases to shading and inundation levels. However, in the case of the other three plants species, the responses varied with a high-water level, with the dry root weight of *C. arborescens, D. zibethinus* and *N. lappaceum* increasing at 30% shading level and decreasing at 70% shading (Table 5).

**Table 5.** Dry weight (DW) of roots of four plant species under treatment combination of shading and inundation.

| Treatment | Dry weight of roots (g) |  |  |  |
| --- | --- | --- | --- | --- |
|  | *C. arborescens* | *S. balangeran* | *D. zibethinus* | *N. lappaceum* |
| Freely drained without shading | 0.75 ± 0.12 c | 0.30 ± 0.07 e | 1.66 ± 0.23 a | 1.20 ± 0.37 b |
| Freely drained with 30% shading | 0.43 ± 0.11 d | 0.24 ± 0.06 e | 0.81 ± 0.30 c | 0.58 ± 0.19 d |
| Freely drained with 70% shading | 0.08 ± 0.03 f | 0.09 ± 0.02 f | 0.30 ± 0.07 e | 0.27 ± 0.18 e |
| GW level 10 cm without shading | 0.55 ± 0.21 d | 0.29 ± 0.03 e | 0.31 ± 0.04 e | 0.50 ± 0.11 d |
| GW level 10 cm with 30% shading | 0.56 ± 0.17 d | 0.21 ± 0.12 ef | 0.60 ± 0.26 cd | 0.70 ± 0.21 c |
| GW level 10 cm with 70% shading | 0.24 ± 0.09 e | 0.10 ± 0.02 ef | 0.37 ± 0.16 e | 0.30 ± 0.08 e |
| GW level zero without shading | 0.76 ± 0.18 c | 0.26 ± 0.01 e | - | 0.32 ± 0.18 e |
| GW level zero with 30% shading | 0.75 ± 0.19 c | 0.19 ± 0.05 ef | 0.34 ± 0.13 e | - |
| GW level zero with 70% shading | 0.14 ± 0.04 ef | 0.07 ± 0.03 f | 0.30 ± 0.86 e | 0.15 ± 0.14 f |
| 1/3 stem height flooding without shading | 0.48 ± 0.20 d | 0.11 ± 0.05 ef | - | 0.05 ± 0.09 f |
| 1/3 stem height flooding with 30% shading | 0.33 ± 0.03 e | 0.12 ± 0.06 ef | 0.25 ± 0.15 ef | - |
| 1/3 stem height flooding with 70% shading | - | 0.10 ± 0.01 ef | - | - |
| Grand mean (S.E.m) | 0.42 (0.55) m | 0.17 (0.55) n | 0.49 (0.55) m | 0.35 (0.55) n |

Note: Blank (-) indicates seedlings' mortality. Means followed by standard deviation. Value followed by different letter is significantly different at $p < 0.05$ . Grand mean is compared among species. S.E.m is standard error of mean.

The root/shoot ratios (RSR) for the four plants species under different treatment combinations is shown in Table 6. The GLM analysis showed that inundation and shading significantly affected the RSR of the tested plant species (S4 Table). The manner in which the four plant species responded to the abiotic stress varied. Three plant species, *C. arborescens, S. balangeran* and *N. lappaceum*, recorded the highest RSR in the control treatment, while *D. zibethinus* has the highest RSR in the treatment with 20 cm inundation and 70% shading (Table 6). The increase in the root-shoot ratio shows that the plant was growing under stress conditions. The RSR is characterized by a biphasic dose-response relationship under stress, typical of hormesis [50]. However, this study found that the plant species responded differently to the various water level and shading treatments. *D. zibethinus* recorded the highest RSR under 30% shading, which indicates a relatively high degree of semi-shade tolerance. *S. balangeran* and *N. lappaceum* recorded the highest RSR under full sunlight (0% shading) at all inundation levels. *C. cratoxylum*, a light-tolerant and wetland species, recorded a response combination on RSR to different shading and water level.

**Table 6.** Root/shoot ratio (RSR) of four plant species under treatment combination of inundation and shading.

| Treatment | Root/shoot ratio at 13 weeks after planting |  |  |  |
| --- | --- | --- | --- | --- |
|  | *C. arborescens* | *S. balangeran* | *D. zibethinus* | *N. lappaceum* |
| Freely drained without shading | 0.93 ± 0.20 a | 0.44 ± 0.07 cd | 0.37 ± 0.08 e | 0.59 ± 0.09 b |
| Freely drained with 30% shading | 0.51 ± 0.03 b | 0.36 ± 0.07 e | 0.40 ± 0.07 d | 0.46 ± 0.10 c |
| Freely drained with 70% shading | 0.23 ± 0.02 g | 0.24 ± 0.07 g | 0.24 ± 0.09 fg | 0.33 ± 0.06 ef |
| GW level 10 cm without shading | 0.65 ± 0.27 b | 0.39 ± 0.03 e | 0.10 ± 0.01 h | 0.40 ± 0.03 d |
| GW level 10 cm with 30% shading | 0.49 ± 0.14 c | 0.23 ± 0.07 g | 0.19 ± 0.10 gh | 0.36 ± 0.04 e |
| GW level 10 cm with 70% shading | 0.34 ± 0.10 def | 0.27 ± 0.03 fg | 0.15 ± 0.03 h | 0.37 ± 0.02 e |
| GW level zero without shading | 0.56 ± 0.07 b | 0.37 ± 0.07 e | - | 0.40 ± 0.004 d |
| GW level zero with 30% shading | 0.48 ± 0.06 c | 0.31 ± 0.04 f | 0.12 ± 0.05 h | - |
| GW level zero with 70% shading | 0.31 ± 0.06 f | 0.20 ± 0.05 g | 0.59 ± 0.46 b | 0.42 ± 0.14 cd |
| 1/3 stem height flooding without shading | 0.38 ± 0.06 e | 0.12 ± 0.05 h | - | 0.08 ± 0.14 h |
| 1/3 stem height flooding with 30% shading | 0.30 ± 0.04 f | 0.10 ± 0.03 h | 0.15 ± 0.08 h | - |
| 1/3 stem height flooding with 70% shading | - | 0.24 ± 0.11 fg | - | 0.12 ± 0.20 h |

Note: Blank (-) indicates seedlings' mortality. Means followed by standard deviation. Value followed by different letter is significantly different at $p < 0.05$ .

Based on the GLM analysis (S4 Table), it can be seen that the interaction between inundation and shading significantly affected the RSR. Single factors related to inundation or shading also affected the RSR. Therefore, we further analyzed the distribution of dry matter growth between shoots and roots in the four plant species studied on the basis of inundation levels, as shown in Fig 3.

**Fig 3.**
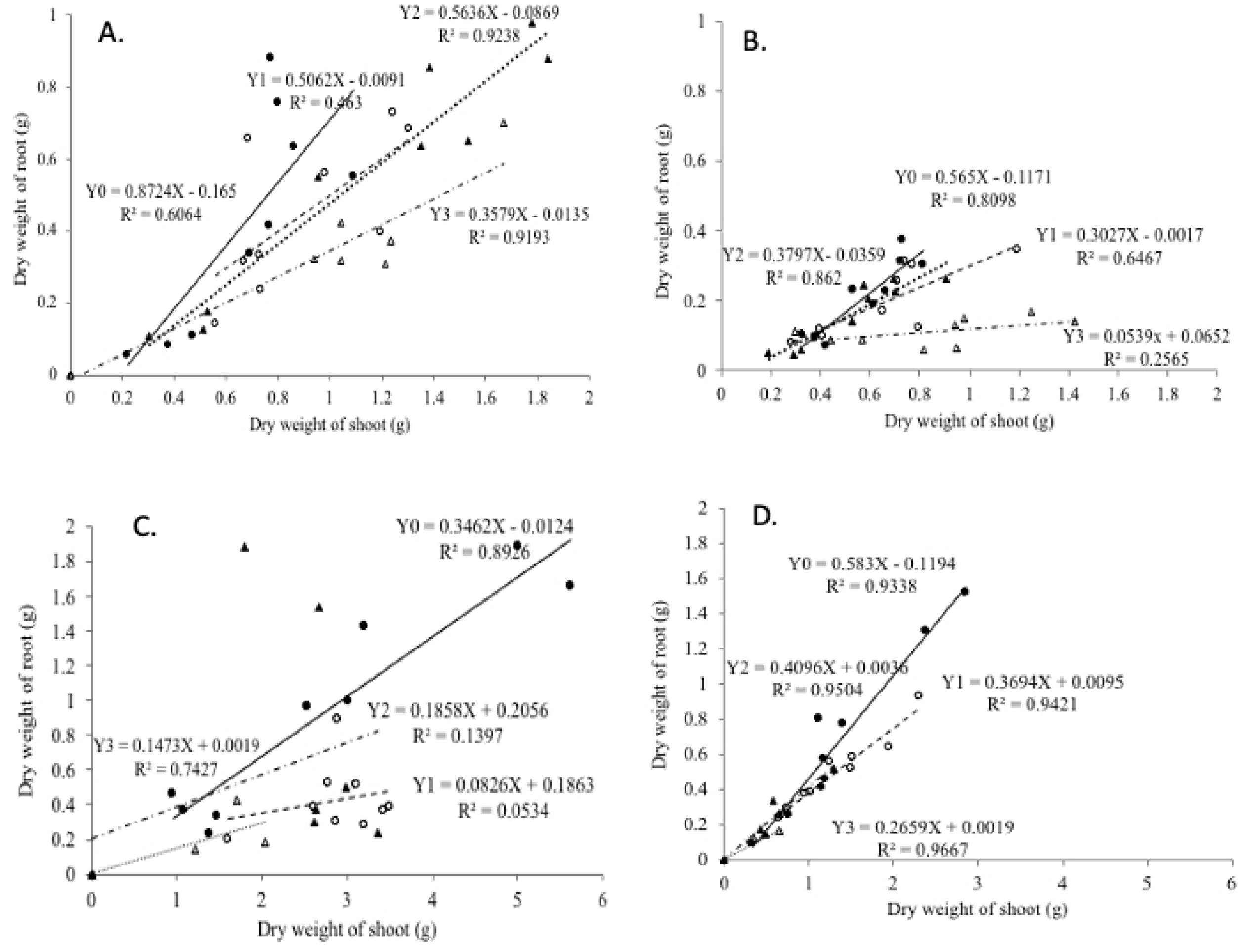
Regression of shoot dry weight on root dry weight in seedlings of 4 plant species at 4 inundation levels: dark circle (Y0) freely drained, light circle (Y1) GW level 10 cm indicates 10 cm inundation, dark triangle (Y2) GW level zero, light triangle (Y3) ⅓ stem height flooding. a) *C. arborescens*, b) *S. balangeran*, c) *D. zibethinus*, d) *N. lappaceum*. Each point indicates the average from 3 samples.

The four plants species displayed a linear relationship between the distribution of dry weight of shoot and root under the 4 inundation levels (Fig 3). The allometric relationship between dry shoot and root weight at the different inundation levels has a different slope and coefficient. This implies that the effect of flooding on the dry shoot and root weight varies between the four tested plant species. The four plants species grew best in non-saturated conditions, although *C. arborescens* and *S. balangeran* are known to grow naturally in peat-swamp forests.

##### Adventitious roots of *C. arborescens* and *S. balangeran*

*C. arborescens* and *S. balangeran* have a relatively high level of adaptability to saturation compared to *D. zibethinus* and *C. arborescens*, with the former able to develop adventitious roots that grow on the stem under the ⅓ stem height flooding conditions. The number of adventitious roots under the ⅓ stem height inundation conditions, under the 3 shading levels, is shown in Fig 4. No adventitious roots developed the cases of *D. zibethinus* and *N. lappaceum*.

**Fig 4.**
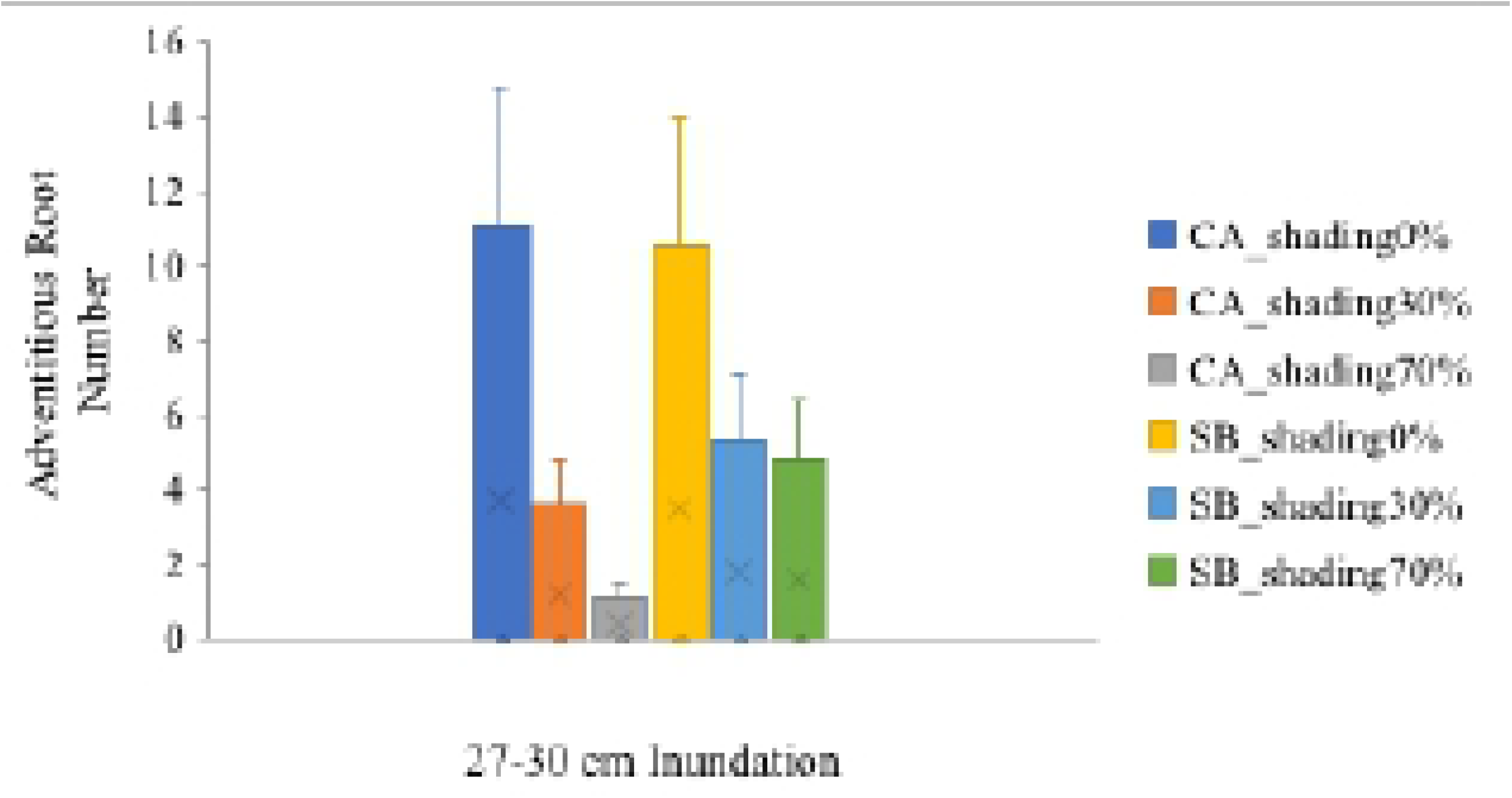
Number of adventitious roots on *C. arborescens* (CA) and *Shorea balangeran* (SB) at 3 levels of shading and ⅓ stem height flooding.

In the treatment involving ⅓ stem height flooding, at all 3 shading levels, the adventitious roots’ primordia were seen in the case of *S. balangeran* in the third week, while the adventitious roots developed in the 5^th^ week. In the case of *C. arborescens*, the adventitious roots developed in the 6^th^ week. The morphology of *S. balangeran*’s adventitious roots differed from those of *C. arborescens*. The adventitious roots in the case of *S. balangeran* persisted in the stem throughout the 13 weeks observation period, while the adventitious roots in the case of *C. arborescens* not persisting, with the number of varying between weeks. The adventitious root formation on the stems in the cases of *C. arborescens* and *S. balangeran* is shown in Fig 5. The anatomy and histology of the adventitious roots are beyond the scope of this study.

**Fig 5.**
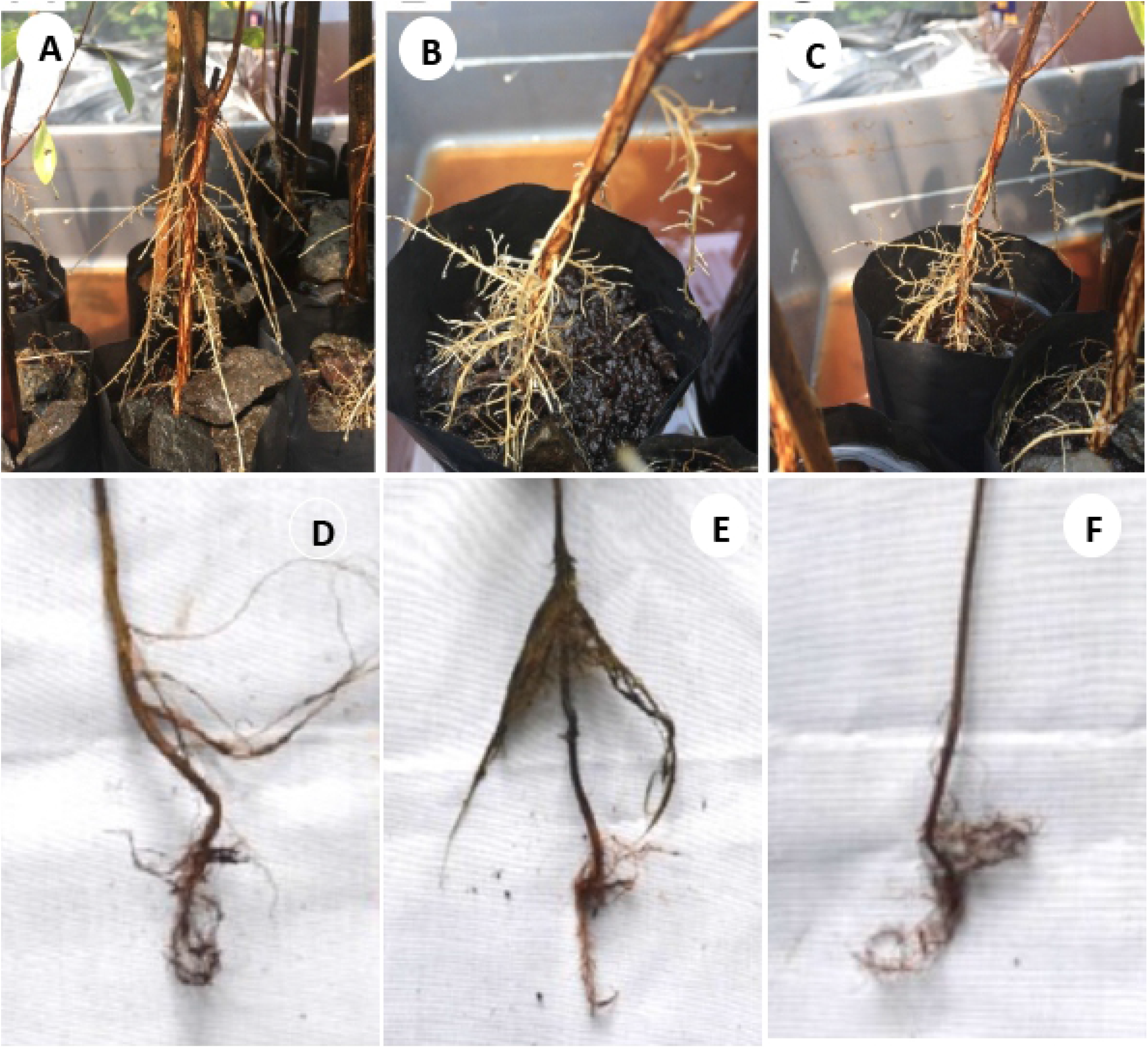
Adventitious root formation of *C. arborescens* (a-c) and *S. balangeran* (d-f) at the ⅓ stem height flooding. (A) & (D) 0% shading; (B) & (E) 30 % shading; (C) & (F) 70% shading.

##### Photosynthesis rate

Treatment combination of species and shading significantly affected the photosynthetic rates of the four treated species (S5 Table). No shading resulted in a relatively higher photosynthesis rate than in the case of the treatments involving 30% and 70% shading, with the photosynthesis rate gradually decreasing with increases to the shading and inundation levels. *D. zibethinus* had the highest photosynthesis rate of all four plant species (Table 7).

**Table 7.** Photosynthesis rate of four plant species under different level of inundation and shading.

| Treatment | Photosynthesis rate ($\mu\text{mol CO}_2 \text{ m}^{-2} \text{ s}^{-1}$) | | | |
| --- | --- | --- | --- | --- |
|  | *C. arborescens* | *S. balangeran* | *D. zibethinus* | *N. lappaceum* |
| Freely drained without shading | 18.2 ± 0.9 bc | 19.3 ± 4.6 b | 24.6 ± 2.2 a | 20.8 ± 2.9 b |
| Freely drained with 30% shading | 9.7 ± 1.0 d | 10.0 ± 1.6 cd | 11.6 ± 0.9 c | 8.3 ± 3.5 d |
| Freely drained with 70% shading | 5.0 ± 1.2 e | 4.8 ± 2.3 ef | 6.1 ± 0.4 e | 2.8 ± 0.07 g |
| GW level 10 cm without shading | 17.9 ± 1.0 bc | 19.6 ± 2.2 b | 24.6 ± 2.3 a | 21.6 ± 3.4 ab |
| GW level 10 cm with 30% shading | 11.0 ± 1.1 c | 9.4 ± 2.6 cd | 10.7 ± 2.6 cd | 8.0 ± 2.0 d |
| GW level 10 cm with 70% shading | 4.4 ± 1.4 ef | 5.7 ± 2.9 e | 5.7 ± 0.2 e | 3.7 ± 0.9 f |
| GW level zero without shading | 17.2 ± 3.3 bc | 19.3 ± 2.7 b | 21.7 ± 1.5 b | 19.0 ± 2.5 b |
| GW level zero with 30% shading | 9.0 ± 1.5 d | 8.0 ± 1.5 d | 8.5 ± 1.3 d | 6.8 ± 2.0 e |
| GW level zero with 70% shading | 3.9 ± 0.8 ef | 4.9 ± 1.4 e | 4.5 ± 0.5 e | 3.8 ± 1.1 fg |
| 1/3 stem height flooding without shading | 16.6 ± 2.4 bc | 18.2 ± 2.9 bc | 18.8 ± 1.1 bc | 11.3 ± 6.0 cd |
| 1/3 stem height flooding with 30% shading | 8.1 ± 1.5 d | 8.29 ± 0.9 d | 9.3 ± 1.3 cd | - |
| 1/3 stem height flooding with 70% shading | 3.4 ± 1.7 f | 3.9 ± 1.6 f | 4.2 ± 1.3 ef | 3.6 ± 1.8 f |

Note: blanks indicate seedling mortality. Mean is followed by standard deviation. Value followed with different letter is significantly different at $p<0.05$ .

## Discussion

### Survival and growth of seedlings

The four tree species responded differently to the treatments given in the experiment. Our study showed that *C. arborescens* and *S. balangeran* seedlings were able to survive and grow in saturated conditions (up to ⅓ of seedling height). *S. balangeran* survived for 13 weeks under stress conditions involving permanent flooding (up to ⅓ of seedlings’ height), under 3 shading levels. However, permanent flooding under 70% shading were not favorable for *C. arborescens* and the two fruit species. *C. arborescens* is a light timber with wood density of 0.61 g cm^-3^ [39], and is categorized as light-demanding [55,56]. *S. balangeran* is a hard-timber with a wood density of 0.86 g cm^-3^ [39]. It has widely distributed habitat and is found in a variety of environments, including seashore, heath forest, peat-swamp forest and lowland forest [39]. Our study showed that under full sunlight (0% shading), *S. balangeran* seedlings grow more slowly than under 30% shading.

The non-peatland tree-species, *N. lappaceum* and *D. zibethinus*, also recorded varying responses to the different shading and inundation treatments. Our study showed that *N. lappaceum* was more tolerant to light than *D. zibethinus* (Table 1, Fig 2). However, seedlings of neither of these two species survived in the high water-level treatments (Table 1, Fig 2). The insufficient O_2_ concentration at the roots probably resulted in decreased root growth which, together with reduced nutrient uptake, resulted in root disfunction that stopped shoot growth [57,58]. (Herzog et al. 2016; Herzog 2017).

The RSR is used as an index to determine the extent of tree growth [51]; it’s indicating the potential of supportive functions relative to the potential of growth functions. Supportive functions consist of water and nutrient uptake, while growth functions consist of light energy harvest [49]. This study showed variations in the RSR recorded between the different species and treatments. *C. arborescens, S. balangeran* and *N. lappaceum* had the highest RSR in the case of the control treatment, except in the case of *D. zibethinus*, for which the highest RSR was recorded in the case of the 20 cm inundation with 70% shading treatment. Variations in RSR are a common stress response driven by the strategies adopted by different plants to cope with stress [49].

#### Adaptability to high water-levels and shading

Sustained high water-level and shading levels affected the growth of all four species of seedlings. High water levels or water logging reduce O_2_ concentration, resulting in hypoxia or anoxia [57]. Flood-tolerant species or peatland species are able to produce aerenchyma on roots and shoots, forming large adventitious root systems with high porosity after logging to enable low-resistance gas transport between plant organs [9,59]. Sustained waterlogging that results in a limited supply of oxygen results in the peatland species, *C. arborescens* and *S. balangeran*, developing adventitious roots, which assist the plant to take up oxygen and transport it into body of the plant [9,57]. By contrast, the two fruit species did not develop adventitious roots and hence did not survive at high waterlogging levels. The adventitious roots were developed only in the case of the ⅓ stem height flooding treatment, at all shading levels. *S. balangeran* had the highest survival rate in the case of all treatments. At the highest waterlogging level, *S. balangeran* developed more adventitious roots than did *C. arborescens*. Visually, the adventitious roots of *S. balangeran* differ from those of *C. arborescens*. Further study on the anatomy and physiology of the treated seedlings involved in this experiment will be reported elsewhere.

The plants’ response ability to flooding stress results from morphological, physiological and metabolic adaptation [60]. Under inundation and heavy shading conditions, the photosynthesis rates in the case of all four plants species was very low, with flooding reducing the photosynthesis rate in the treatments of all four species (Table 7). This may be caused by the closure of stomata at high water-levels, which restricts internal CO_2_ concentrations and results in negative feedback due to carbohydrate accumulation [57]. Moreover, light intensity levels under heavy shading conditions are also limited. These treatment combinations resulted in inhibited root growth and nutrient uptake. Waterlogged soil resulted in the roots of the fruit tree species (*D. zibethinus* and *N. lappaceum*) having a low biomass. However, the adventitious roots that grow on the stems of *C. arborescens* and *S. balangeran* enable some nutrient uptake and respiration to occur.

### Further research needs and its application

Our study shows the manner in which the rates of survival, growth and adventitious root formation for four commonly planted tropical peatland species vary under different inundation and shading conditions. These findings can be applied to enable planning processes related to restoration initiatives in Indonesia, with different restoration sites having different hydrological and shading conditions. As reported earlier, some agroforestry systems located in peatlands in Central Kalimantan have deep water levels. For instance, the ground water level in an agroforest that includes N. *lappaceum* and *Dyera polyphylla* tree species in Tumbang Nusa village of Central Kalimantan province in 2014 fluctuated from between −20 to −140 cm [28], whereas the ground water level in another agroforest farm that included fruit tree species in Henda village, Central Kalimantan, ranged from −8 to −60 cm in 2019 [44]. However, flooding in the area was not reported. Therefore, it can be concluded that *D. zibethinus* and *N. lappaceum* can grow well on peatlands in Central Kalimantan.

According to the Government Regulation No. 57, 2016, peat ecosystems that are located between two rivers, or between river and sea, and /or on swamp are referred to as a Peatland Hydrological Unit (PHU). Peatland with a peat-depth of less than 3 m is categorized as having a cultivation function, while peat with peat-depth of greater than 3m is categorized as having a protected function. A ground water level of −40 cm is mandatory in cases of peatland with a cultivation function, while in the case of peatlands with a protected function, the water level should be close to the ground [17]. In rewetted peatlands, the planting of peatland tree species, such as *S. balangeran, C. arborescens. N. lappaceum*, is recommended, as these can survive and grow at GWL 10 cm. These species could also be planted in rewetted peatlands in agroforestry or mixed planting systems. Light-tolerant species, such as *C. arborescens*, could be planted prior to planting *S. balangeran* and *N. lappaceum*. It is necessary to apply land preparation and management processes, such as mounding, prior to the planting of flooding-intolerant tree species (such as *D. zibethinus*). The use of a combination of timber and fruit tree species would provide ecological benefit for the ecosystem and economic benefit for the communities.

## Acknowledgement

This study received financial support from the Partnership for Enhanced Engagemnet in Research (PEER) USAID, Grant Number 2000008728, under a research project collaboration (“Developing Biodiverse Agroforests of Rewetted Peatlands”) between the World Agroforestry Center and the Forest Research and Development Center (FRDC). The nursery experiment was supported by the Faculty of Forestry and Environment at the IPB University, through a collaboration between the FRDC of the Ministry of Environment and Forestry and IPB University.

## Contributions of Authors

HLT, HSN, and Is together designed the experiment and supervised and coached four undergraduate students at IPB University. DAH, HA, NH, MP conducted the experiment and analyzed the initial data. HLT re-analyzed data compilation; HLT and HSN wrote the first and improved manuscript; MvN, RC and RK provided guidelines and were also involved in writing the improved manuscript. All authors read and agreed with the MN findings presented in this manuscript.

